# Genome-wide, time-sensitive interrogation of the heat shock response under diverse stressors via ReporterSeq

**DOI:** 10.1101/2020.03.29.014845

**Authors:** Brian D. Alford, Gregory Valiant, Onn Brandman

## Abstract

Interrogating cellular stress response pathways is challenging because of the complexity of regulatory mechanisms and response dynamics, which can vary with both time and the type of stress. We developed a reverse genetic method called ReporterSeq to comprehensively identify genes regulating a stress-induced transcription factor under multiple conditions in a time-resolved manner. ReporterSeq links RNA-encoded barcode levels to pathway-specific output under genetic perturbations, allowing pooled pathway activity measurements via DNA sequencing alone and without cell enrichment or single cell isolation. Here, we used ReporterSeq to identify regulators of the heat shock response (HSR), a conserved, poorly understood transcriptional program that protects cells from proteotoxicity and is misregulated in disease. We measured genome-wide HSR regulation in budding yeast across thirteen stress conditions, uncovering novel stress-specific, time-specific, and constitutive regulators. ReporterSeq can assess the genetic regulators of any transcriptional pathway with the scale of pooled genetic screens and the precision of pathway-specific readouts.

## Introduction

The heat shock response (HSR) is a conserved transcriptional response that protects cells from cytoplasmic proteotoxicity (Lindquist, 1986; Morimoto, 2011). The HSR is driven by the transcription factor Hsf1, a trimeric protein which promotes expression of chaperones and other cellular components that refold misfolded proteins or target them for degradation (Akerfelt et al., 2010). A diverse array of stressors activate the HSR, including heat, ethanol, oxidative stressors, and amino acid analogs (Morano et al., 2012). In these conditions, HSR activity can follow diverse trajectories, with both transient or prolonged activity observed (Sorger, 1990). Furthermore, aberrant HSR activation has been linked to neurodegeneration (too little HSR activation; (Campanella et al., 2018)) and cancer (too much HSR activation; (Whitesell and Lindquist, 2009).

The complexity of the HSR has led to a multitude of models for how the HSR is regulated (Anckar and Sistonen, 2011). These include negative regulation in which the chaperones Hsp70 (Krakowiak et al., 2018; Zheng et al., 2016), Hsp90(Ali et al., 1998; Zou et al., 1998), and Hsp60(Neef et al., 2014) bind and inhibit Hsf1 directly. Signaling pathways have also been demonstrated to modulate Hsf1 activity through phosphorylation by kinases such as Snf1, Yak1, and Rim15 (Hahn and Thiele, 2004; Lee et al., 2013, 2008). Additionally, post-transcriptional models have been proposed, such as HSR-regulated mRNA half life (Heikkinen, 2003) or regulation of the translation efficiency of heat shock mRNAs (Zid and O’Shea, 2014). Yet the extent to which these and other mechanisms drive the HSR under diverse stressors is poorly understood, limiting our ability to understand and remedy the HSR in disease states (Neckers and Workman, 2012).

To understand the molecular mechanisms driving the HSR over time in diverse stress conditions, it is critical to identify the genes involved. Reverse genetic studies of the HSR have been useful in discovering both new protein quality control machinery in the cell (Brandman et al., 2012), as well as new methods of regulating the HSR (Raychaudhuri et al., 2014). However, existing reverse genetic approaches to measure specific pathway activity require measuring the effect of each gene separately or enriching cells according to pathway activity, making it impractical to test multiple environmental conditions or make time-resolved measurements.

To address this methodological deficit and knowledge gap, we developed ReporterSeq, a pooled, genome wide screening technology that can measure the effect of genetic perturbations with the scale of pooled genetic screens and the precision of pathway-specific readouts (e.g. qPCR, fluorescence, luciferase reporters). ReporterSeq measures pathway activity under a specific perturbation through levels of a corresponding barcode. Thus, ReporterSeq does not require cell sorting and instead relies only upon measuring barcode frequencies at a single genetic locus of a pooled sample. This allows dozens of samples to be collected on the same day, prepared for sequencing, and then read out in a single high throughput sequencing run. Because ReporterSeq uses RNA levels as a direct readout of transcriptional activity, pathway activity can be measured even in conditions in which protein synthesis is compromised.

We performed ReporterSeq in the budding yeast, *Saccharomyces cerevisiae*, under basal growth conditions and thirteen stress conditions to identify general and stress-specific regulators of the HSR. This revealed the roles of known regulators, like the Hsp70 chaperone system and Snf1 kinase complex, as well as novel regulators, including Gcn3 and Asc1, in responding to diverse stressors. A time course of HSR activation in response to heat stress revealed distinct regulatory mechanisms in the early and late stages of heat stress. We further investigated how loss of Gcn3 regulates the HSR and found that, contrary to its canonical role in stress-dependent inhibition of translation, Gcn3 is required for efficient translation of the heat shock reporter transcript under arsenite stress. In addition to elucidating HSR regulation, our work demonstrates that ReporterSeq is a generally applicable tool to dissect the genetic basis for the activity of any transcription factor in a quantitative, scalable, and time-resolved manner.

## Results

### A pooled strategy to measure expression of a specific gene under genome-wide, CRISPR perturbations without cell enrichment

ReporterSeq measures how endogenous genes affect the expression of an exogenous reporter gene. It accomplishes this through pairing a genetic perturbation with barcode levels from a pathway-specific readout (Figure 1). Though ReporterSeq can be implemented with any encodable genetic perturbation (e.g. RNAi, CRISPR knockouts), we used CRISPRi (Gilbert et al., 2013) as a perturbation to lower expression of each gene and paired it with a barcoded mRNA encoding GFP downstream of an Hsf1-responsive synthetic promoter built upon a “crippled” CYC1 promoter sequence derived from a 225 nucleotide fragment of the CYC1 promoter, similar to that used in a fluorescence based HSR screen(Brandman et al., 2012; Guarente and Mason, 1983). To maximize dCas9-Mxi1 (a transcription inhibitor) and sgRNA expression, we expressed dCas9-Mxi1 and the ReporterSeq cassette on two separate high copy plasmids that were coexpressed in each cell. We verified that the genetic knockdowns could modulate the heat shock reporter by measuring the effects of known HSR regulators (Brandman et al., 2012) on reporter output (Figure S1A). We were therefore confident that CRISPRi knockdowns were effective in our system and capable of altering HSR reporter activity.

**Figure 1.**
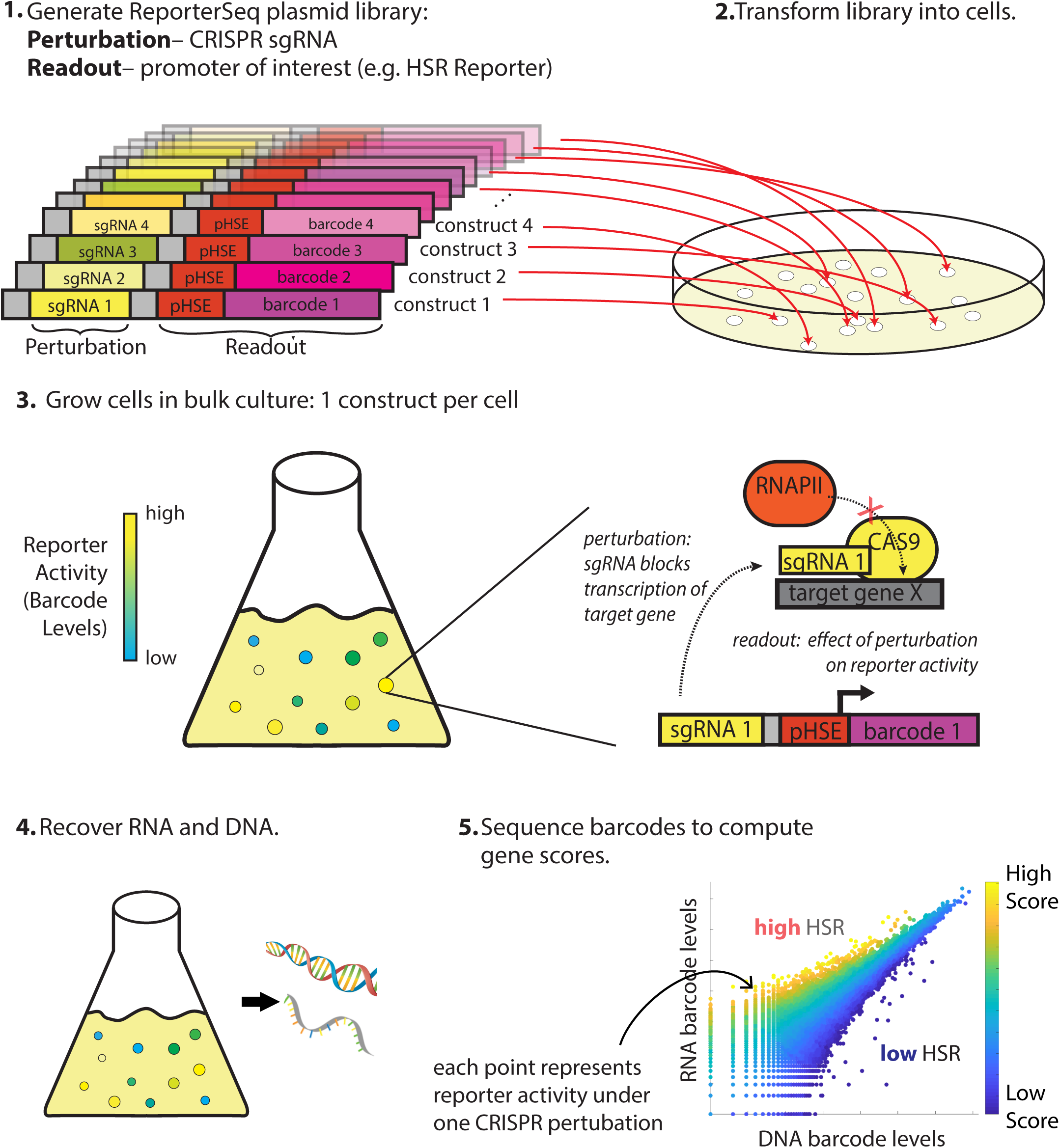
Overview of ReporterSeq method. Example given for measuring the heat shock response (HSR) using a heat shock element (HSE)-driven promoter and CRISPRi to perturb gene expression. ReporterSeq measures the effect of genetic perturbations with the throughput of pooled screens and the precision of pathway-specific, transcriptional readouts.

The first step of ReporterSeq is to synthesize a diverse library of constructs, each containing one sgRNA and one barcode driven by a promoter of interest. The library is then introduced into cells such that each cell contains one construct. Cells are grown in the appropriate conditions and then harvested at the desired timepoints. RNA and DNA are harvested from the cells and the counts of each barcode are tallied using deep sequencing. The RNA counts for a given barcode are proportional to the total transcriptional output of all cells containing that barcode, while the DNA barcode counts are proportional to cell numbers. Thus, the RNA/DNA ratio of a given barcode reflects the activity of the reporter under the influence of the specific CRISPR knockdown corresponding to that barcode. This ratio is used to identify the effect of a given knockdown on the HSR in a single condition. In situations where the same pool of yeast is divided and exposed to multiple conditions, RNA counts between these samples can be directly compared to compute gene-stressor interactions (discussed below).

Our library targeted each gene in the genome, including genes producing non-coding RNAs, with up to twelve sgRNAs and paired multiple random barcodes with each sgRNA. Multiple sgRNAs per gene were necessary due to variability in efficacy of each knockdown and to minimize the influence of off-target effects from any one guide RNA. Using multiple barcodes for each sgRNA minimized the effects of barcode bias on stability of the reporter mRNA. We cloned a library of approximately 1 million random barcode-sgRNA pairs and performed paired-end high-throughput sequencing on the library to identify these pairs. Paired end sequencing revealed that most genes were targeted in the library by all twelve designed sgRNAs (Figure S1B) Additionally, there was a wide distribution of barcodes associated with each sgRNA, with the mean representation being approximately fourteen barcodes per sgRNA (Figure S1C). Thus, the plasmid library was poised to measure the effect of each genetic knockdown on the output of the HSR reporter.

### ReporterSeq reveals regulators of the HSR in basal conditions

We first used the ReporterSeq library to measure the effect of each genetic knockdown on the HSR reporter in untreated, log-phase yeast. The full genome library was transformed into >100,000 cells, resulting in a subset of the plasmid library expression in cells. RNA and DNA barcodes were sequenced and matched to their corresponding sgRNAs, revealing that 91% of genes were represented by 6 or more sgRNAs (Figure S1D) and 74% of sgRNAs were represented by 2 or more barcodes (Figure S1E) where both RNA and DNA counts were above 10. To test if the RNA/DNA ratio of barcodes reflected activation of the HSR under a specific perturbation, we assessed the effect of two predicted regulators of our synthetic HSR reporter (Figure 2A). sgRNAs targeting *SSA2*, an Hsp70 chaperone that binds Hsf1 and inhibits the HSR (Krakowiak et al., 2018; Zheng et al., 2016), resulted in a higher than average RNA/DNA ratio as expected. Conversely, sgRNAs targeting *CYC1*, the gene from which our synthetic reporter was derived and shares significant homology with, lowered the RNA/DNA ratio compared to average. Thus, the DNA/RNA ratio for a single barcode reflects the HSR reporter’s activity under a specific genomic perturbation.

**Figure 2.**
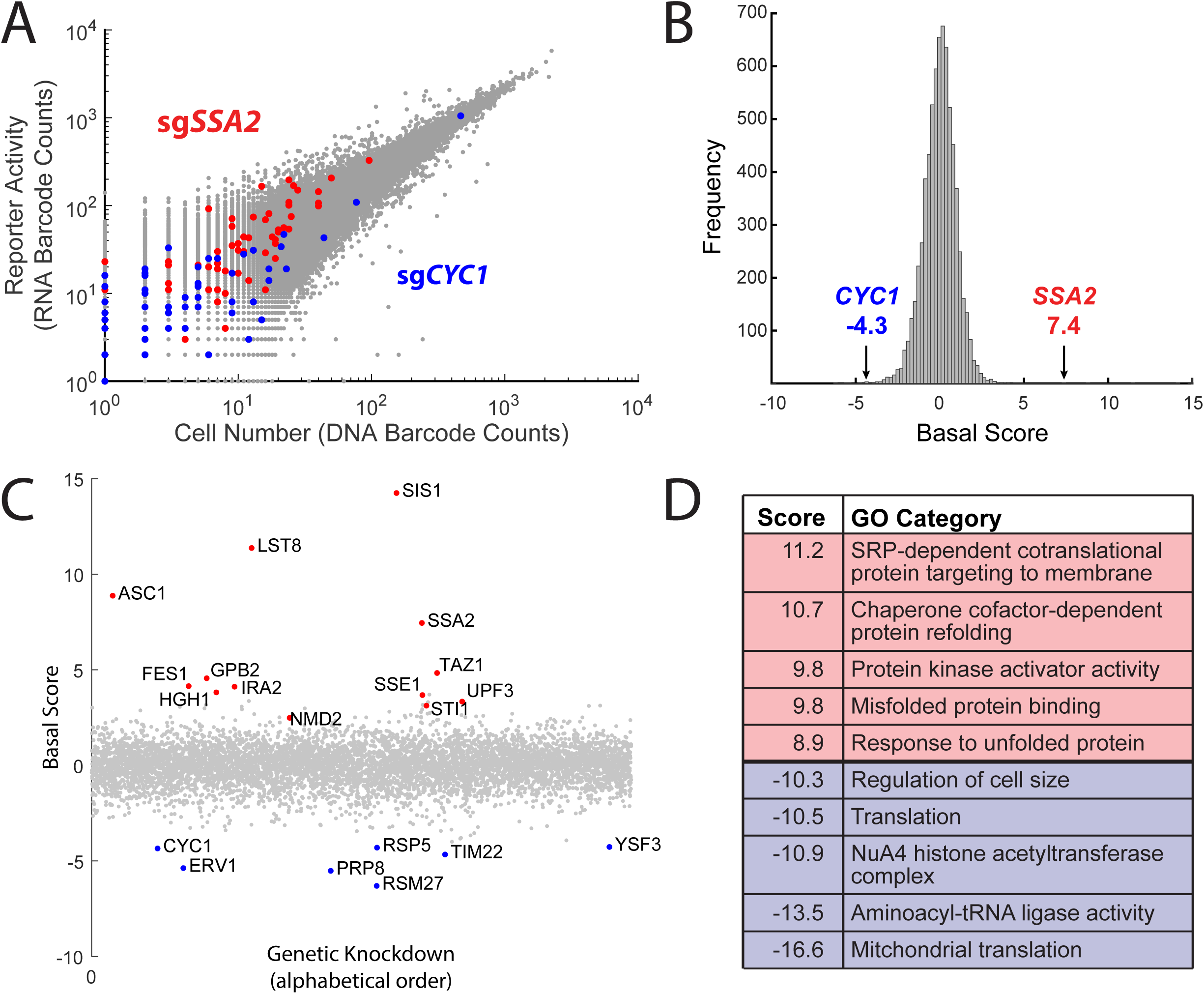
ReporterSeq reveals genome wide basal regulators of the heat shock response. A.) Comparison of DNA and RNA barcode counts from wildtype, untreated yeast. Red dots indicate barcodes that correspond to sgRNAs targeting the Hsp70 chaperone, *SSA2*. Blue dots indicate barcodes that correspond to sgRNAs targeting *CYC1*,a gene which shares significant homology with the Hsf1-driven synthetic reporter. B.) Histogram of the basal scores for sgRNAs targeting each gene. Scores for *SSA2* and *CYC1* gene targets are indicated. C.) Plot of the basal scores for every genetic knockdown. Outliers and genes mentioned in the text are labeled. D.) Table of the gene ontology (GO) categories with the 5 highest and 5 lowest scores in the basal screen.

To facilitate comparisons between many data points, we developed a scoring method that combined the information from the multiple barcodes and corresponding sgRNAs which target a single gene into a single score, ***s*** (see methods). Briefly, the scoring metric determined the extent to which each barcode was an empirically-tested outlier relative to other barcodes with similar levels of representation in the data. The magnitude of the barcode score indicates the confidence that the gene affects the HSR, while the sign of the score indicates the direction of the effect on the HSR (positive scores indicate that the knockdown increases the HSR, while negative scores indicate the knockdown decreases the HSR). This strategy assumes that most perturbations do not affect the HSR, and will thus have a score near 0. The score for each gene was tallied as the average barcode score over all sgRNAs and barcodes. Scores were then scaled, such that the magnitude of each score represents the number of standard deviations from the mean (Z-score). *SSA2* knockdown, for example, had the fourth highest score at 7.4, indicating that the knockdown of *SSA2* likely increased reporter activity, with the confidence that this occurred being 7.4 standard deviations from the mean (Figure 2B). *CYC1*, conversely, had the 5th lowest score of -4.3, meaning that it likely decreased the reporter activity with less though still substantial confidence (all gene scores are provided in table S1). Our scoring metric thus quantifies the confidence that genes affect reporter activity.

Strong hits were reproducible while weak hits were noisy (duplicates with corrcoef .27 shown in Figure S2; addressed in discussion). To reduce noise, we averaged scores from 4 biological replicates of basal Hsf1 activation (Figure 2C). Lowering expression of the chaperones or cochaperones *SSA2, SIS1* (***s***=14.2), *FES1* (***s***=4.1), *SSE1* (***s***=3.7), *HGH1* (***s***=3.8), and *STI1* (***s***=3.1) strongly activated the HSR. Indeed, the Hsp40 *SIS1* was the strongest basal suppressor of the HSR. Additional strong HSR suppressors include the negative regulators of the Ras/cAMP pathway *GPB2* (***s***=4.6) and *IRA2* (***s***=4.1), the mitochondrial lyso-phosphatidylcholine acyltransferase *TAZ1* (***s***=4.8), the nonsense-mediated decay regulators *UPF3* (***s***=3.3) and NMD2 (***s***=2.5) and the small ribosomal protein Asc1 (***s***=8.9). Lst8 (***s***=11.3) was also a strong hit in the screen, likely because it is adjacent to *SIS1* in the genome and thus silenced by the same sgRNAs (Pincus et al., 2018). Our implementation of ReporterSeq cannot distinguish between adjacent genes that may be targeted by the same sgRNAs, a general shortcoming of CRISPRi. In addition to *CYC1*, several mitochondrial genes (*ERV1* ***s***=-5.4, *RSM27* ***s***=-6.3, and *TIM22* ***s***=-4.7) strongly lowered the HSR when impaired, consistent with previous observations (Brandman et al., 2012) as well as the E3 ubiquitin ligase *RSP5* (***s***=-4.3), which is required to maintain Hsf1 levels (Haitani et al., 2006; Haitani and Takagi, 2008).

In order to identify processes that regulate the HSR, we analyzed which gene ontology (GO) categories caused high or low activation of the heat shock response. Each GO category is a curated set of genes corresponding to a specific cellular function (Ashburner et al., 2000; The Gene Ontology Consortium, 2019). We assigned each GO category a score based on the Student’s t-test between the scores of all genes and those of the specific category (Table S2). As with the interaction score, the sign indicates the direction of the effect of the knockdown on the reporter activity, and the magnitude of the score indicates the confidence. Many of the GO category scores followed trends largely expected from previous studies, with chaperone, protein folding, and protein quality control amongst the most highly scoring GO categories (Figure 2D). The lowest scoring categories included mitochondria and translation, consistent with results of a previous HSR screen (Brandman et al., 2012).

### Interactions between heat stress and genes that regulate the HSR

To determine which genes affect the cell’s ability to activate the HSR in response to heat stress, we sought to quantify the interaction between the effects of heat stress and those of each genetic knockdown (a “stressor-gene” interaction) on the output of the HSR reporter (Figure 3A), similar to how genetic interactions (“gene-gene interactions) are conceptualized (Collins et al., 2007; Tong et al., 2001). Stressor-gene interactions are based on a comparison of the reporter output of the combined condition relative to that of the knockdown and stress independently. A stressor-gene interaction score of zero denotes that knockdown of the gene and the stressor itself have independent effects on the HSR. A positive score denotes that the combination of gene and stress resulted in a superadditive effect on the HSR (an “aggravating” interaction), while a negative score denotes that the combination results in a subadditive effect on the HSR (a “blocking” interaction). A positive or negative interaction score suggests that a stressor and gene are related in the context of the HSR. Traditional methods to compute interaction scores require three measurements: the phenotype of each individual perturbation and the phenotype of the double mutant. By assuming that 1) the majority of genetic knockdowns do not interact with the stressor and 2) short stressor treatments that impair cell growth (relative to the ∼2.5 hour doubling time of unstressed yeast in the growth conditions used) do not significantly alter cell counts for a specific CRISPRi perturbation, we computed stressor-gene interactions by comparing RNA levels between stressed and unstressed conditions from the same yeast sample using the same scoring metric as with the untreated RNA/DNA ratios. This allowed us to compute the stressor-gene interactions without calculating the absolute induction of the gene (its RNA/DNA ratio), reducing noise arising from DNA normalization and increasing robustness by performing more comparisons than traditional methods.

**Figure 3:**
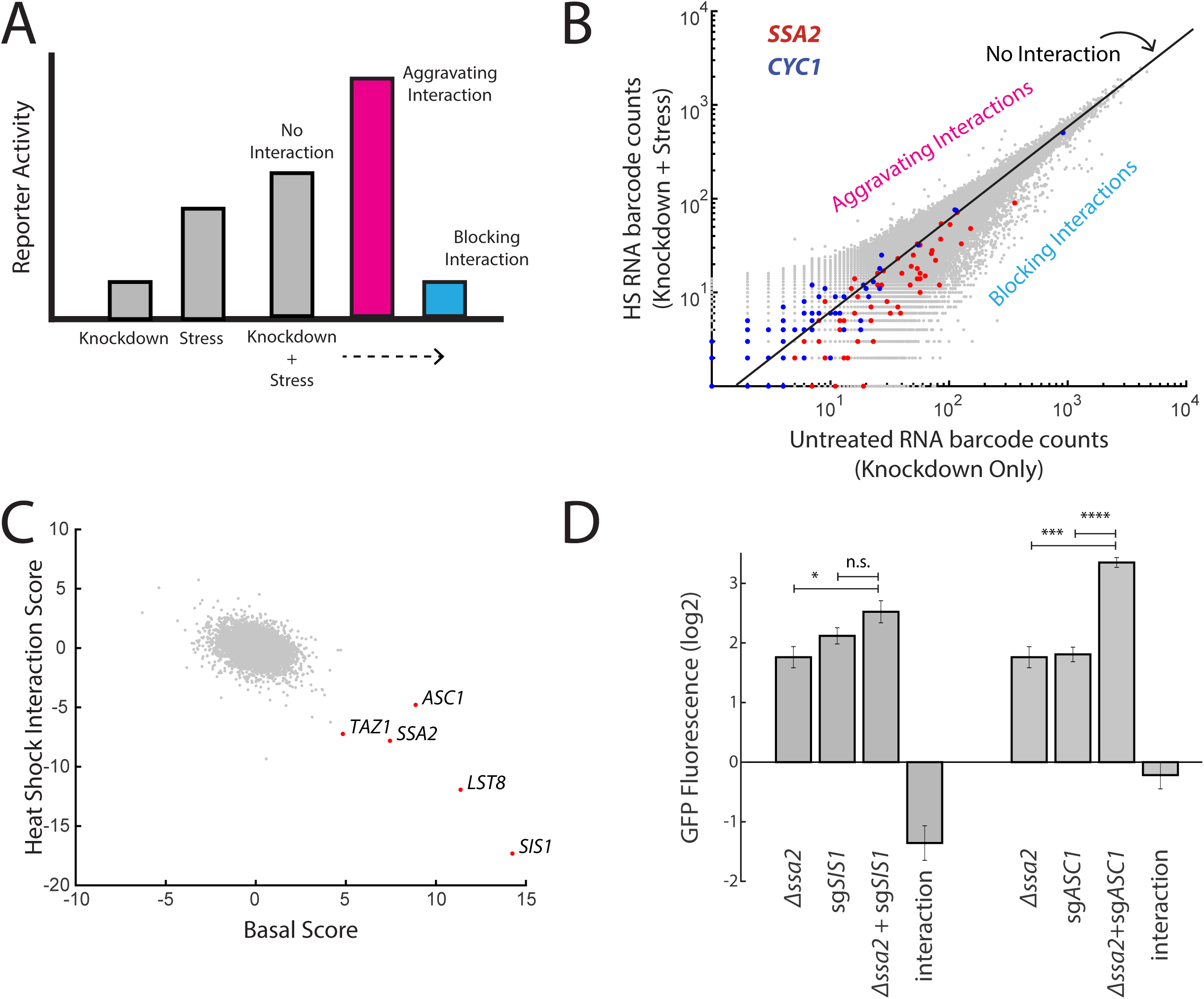
ReporterSeq reveals how genes interact with heat stress. A.) A schematic of gene-stressor interactions. The combination of a gene perturbation and stressor can be additive (no interaction, suggesting independence), higher than expected (an aggravating interaction), or lower than expected (a blocking interaction). B.) RNA barcode counts of untreated yeast compared to those of heat-stressed yeast from the same sample. No interaction, aggravating, and blocking interactions are indicated. Red dots indicate barcodes that correspond to sgRNAs targeting *SSA2*. Blue dots indicate barcodes that correspond to sgRNAs targeting *CYC1*. C.) Basal score for each gene versus the heat shock interaction score for each gene. Genes with high basal score are labeled. D.) Genetic interactions for *Δssa2*-sg*SIS1* and *Δssa2*-sg*ASC1* gene pairs. GFP fluorescence of the HSR reporter in the *ssa2* knockout alone, an sgRNA alone, and both perturbations together are displayed, each relative to wildtype cells containing the reporter. The genetic interaction is the observed combined log2 reporter activity minus the reporter activity in each individual genetic perturbation. Error bars are standard errors of at least 3 replicates. P-values are indicated for select pairs of conditions based on the following key: n.s.-p>0.05; *-p<0.05; **-p<0.01; ***-p<0.001; ****-p<0.0001.

We then examined which genes showed strong interactions with a heat stress of 20 minutes at 39^°^C. As an example of the interaction scores in practice, we first examined *SSA2* and *CYC1. SSA2* had a blocking interaction with heat (***s***=-7.8) whereas *CYC1* had an aggravating interaction with heat stress (***s***=2.3; Figure 3B). This suggests that heat stress and *SIS1* depletion share a pathway of HSR activation and that targeting both the *CYC1* gene as well as the HSR reporter directly (because the reporter is built upon the *CYC1* promoter sequence) do not reduce its inducibility by heat. Indeed, induction was higher than expected in knockdown of *CYC1* (a positive interaction) and, generally, heat stress interaction score and the basal knockdown score were negatively correlated (*r*=-0.40), suggesting that knockdowns basaly increasing the HSR generally blocked HSR activation in heat and those decreasing basal HSR resulted in a bigger HSR increase during heat stress.

One knockdown target that blocked the response to heat less than would be predicted based on the general trend for strong HSR inducers was *ASC1* (Figure 3C). This suggests that activation of the HSR by *ASC1* may be through mechanisms independent of the other genes that strongly activated the HSR, such as *SIS1* and *SSA2*. To test this, we measured the genetic interaction between combined genetic perturbations (deletions combined with CRISPRi) to determine if these genes belonged in a shared or independent pathway of activating the HSR (Figure 3D). As expected, *SIS1* (encoding a cochaperone of Hsp70) and *SSA2* (encoding an Hsp70) had strong blocking interactions with each other, as the combination of the two perturbations was 65% of the expected GFP levels (quantified as log GFP levels). By contrast, ASC1 was independent of SSA2, with the combination of genetic perturbations having an almost completely additive effect on the reporter (GFP levels 94% of expected levels). This suggests that limiting Asc1 levels may activate the HSR through a separate mechanism than limiting Hsp70 levels.

### Distinct genes regulate the early and late phases of the HSR

The pooled format of ReporterSeq and very short sample collection time allowed us to measure the effect of each knockdown on the HSR in a time-resolved way with a precision that genome-wide approaches generally cannot achieve. We collected cells at 10, 20, 40, and 120 minutes after transition to 39°C in duplicate and isolated RNA to calculate stressor-gene interactions each time point (Figure 4A). Most genes had similar interaction scores for the first and last timepoints. For example, knocking down *SIS1* or *TPS1* blocked the HSR throughout the time course (Figure 4B). Tps1 is required for synthesis of trehalose, and has been demonstrated to be required for induction of the HSR under heat shock (Conlin and Nelson, 2007). Other interactions, however, greatly depended on the length of heat shock. Knockdown of the Ras gene, *RAS2*, caused an aggravation of the HSR at 10 and 20 minutes, had no effect at 40 min, and then subsequently blocked the HSR at 120 min. Knockdown of HSF1 had a strong blocking effect at 10 minutes that tapered to near zero by 40 minutes. Conversely, knockdown of *HSP42, BTN2* or *OPI10* had relatively little effect until 40 and 120 minutes, at which point the HSR became hyper-induced. These genes have been reported to be critical for survival in heat shocked cells, with Hsp42 and Btn2 sequestering toxic proteins(Haslbeck et al., 2004; Malinovska et al., 2012; Specht et al., 2011) and the human and Schizosaccharomyces pombe orthologs of Opi10 (“Hikeshi”) facilitating the stress-induced import of Hsp70 into the nucleus(Kose et al., 2012; Song et al., 2015). The delay in the effects of these factors likely represents the time in which cellular damage accumulates under heat shock. Curiously, the nearly identical HSP70 paralogs SSA1 and SSA2 had strikingly different interaction profiles over time, with SSA2 having strong blocking interactions with all time points, and SSA1 mostly not interacting with heat stress but having aggravating interactions at 40 and 120 minutes. These distinct profiles suggest differential regulation of the HSR by SSA1 and SSA2 over prolonged heat stress.

**Figure 4.**
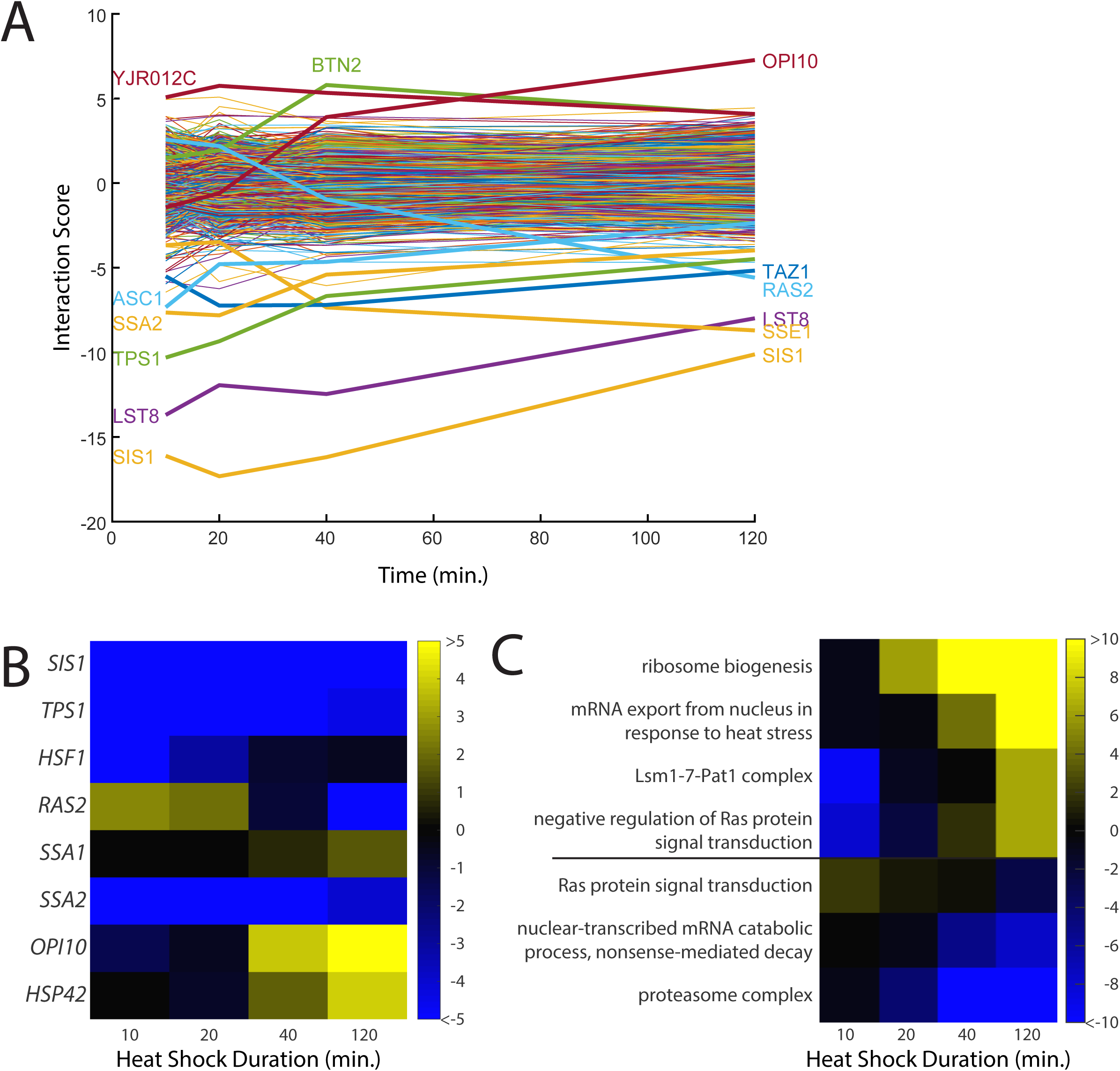
Time-resolved interactions between genetic knockdowns and heat stress. A.) Time course of interaction scores for each gene with 39°C heat shock. Time points were measured at 10, 20, 40, and 120 minutes. B.) Heat map of interaction scores for selected genes in the screen. C.) Heat map of GO category scores for selected categories with differential effects in early and late time points.

To identify cellular processes that affected the early and late responses to heat shock differently, we analyzed which GO categories had differential effects over the time course (Figure 4C; Table S2). Like the aforementioned GO scores, these GO scores were calculated based on the Student’s t test between the scores of all genes and those of the specific category. Knockdown of genes that are a part of the heat shock response, such as “mRNA export from nucleus in response to heat stress”, generally exacerbated the heat shock response at later time points, suggesting that if the response to heat is impaired, the stress response will be prolonged. Another mRNA regulator, the Lsm1-7-PatI complex, which degrades some mRNAs, also caused a stronger late phase heat shock response. Additionally, genes involved in Ras signal transduction followed the same trend as *RAS2* and were unusually high at early time points but low at later ones. Conversely, negative regulators of Ras showed the opposite trend. The lowest interaction scores found in later time points were proteasomal genes, while the highest scores were for ribosome biogenesis genes. ReporterSeq can thus identify cellular components that are critical for different phases of a stress response.

### ReporterSeq reveals stress-specific HSR regulators

We next performed ReporterSeq on twelve additional stress conditions, including nutrient deprivation and toxins (Table S3). These stressors affect cells through a variety of mechanisms and cause varying degrees of HSR reporter activation as measured by qPCR (Figure 5A). We applied these stressors for between 20 minutes and 3 hours. As expected, the pleiotropic proteotoxic stressors and amino acid analogs known to cause accumulation of misfolded protein (Jacobson et al., 2012; Trotter et al., 2002; Weids et al., 2016) robustly activated the HSR. The other stressors, including amino acid starvation, glucose starvation, and the ER stressors DTT and tunicamycin, had milder effects on the HSR. We calculated interaction scores for each gene in each stress condition (Figure 5B). Some of the stressors shared the same strong genetic interactions that we observed under heat shock conditions. For example, all stressors, with the exception of the proteasome inhibitor bortezomib, had a negative interaction score with the SIS1 and SSA2 knockdowns.

**Figure 5.**
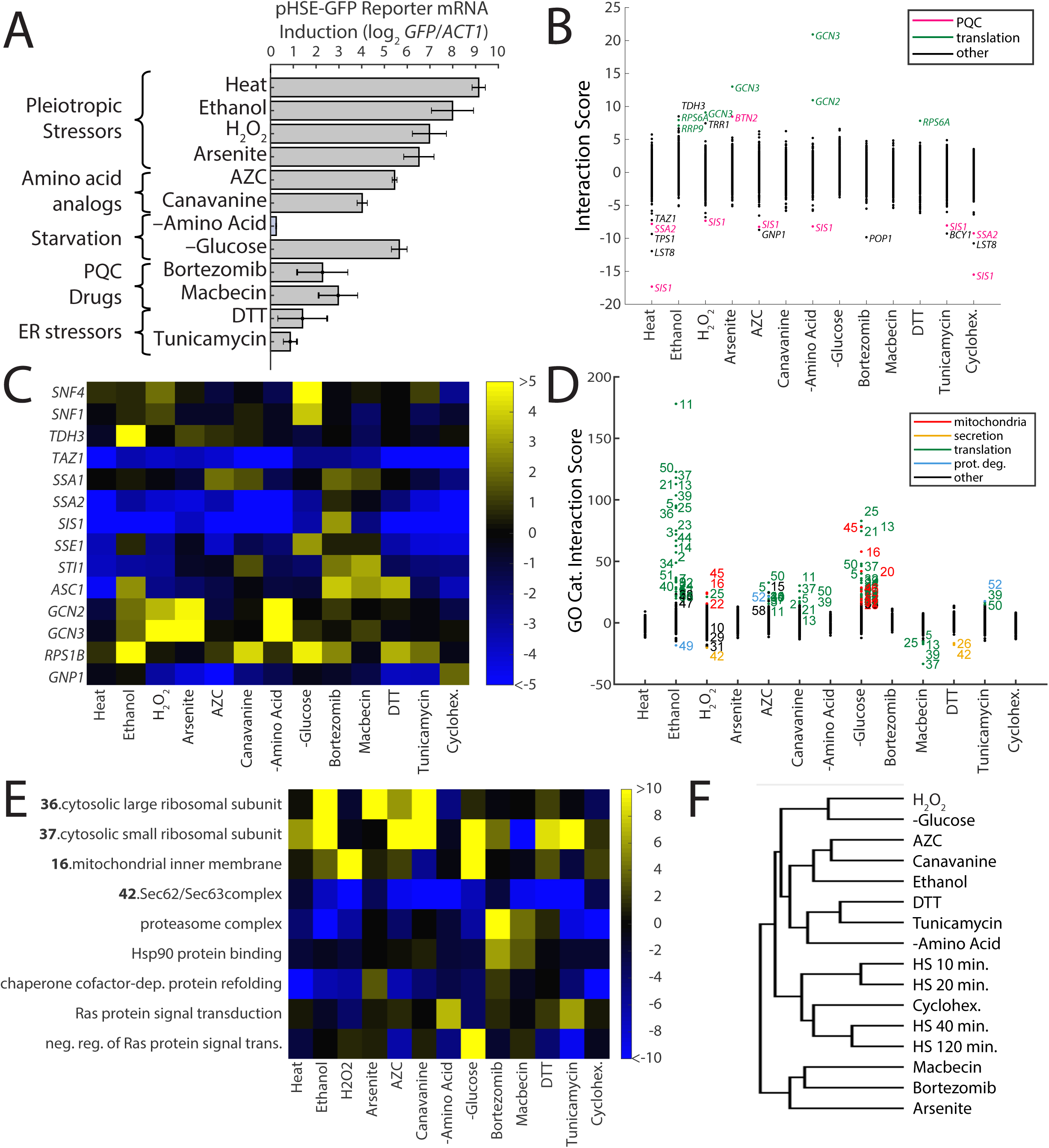
ReporterSeq reveals the diversity of responses to different stress conditions. A.) HSR reporter mRNA induction based as measured by qPCR in 13 stressors (see table S3 for details of treatments). mRNA levels are relative to *ACT1* mRNA levels, and the results are normalized such that untreated yeast have an induction of 0. Error bars are standard errors of at least 3 replicates. B.) Interaction scores for each gene under each stressor. Strong outliers are labeled and colored based on annotated function. C.) Heat map of interaction scores for select genes with each stressor. D.) GO category scores for each gene under each stressor. Category ID number key is provided in table S4. Outlier categories are colored based on function. E.) Heat map of GO category scores for selected GO categories and each stressor. F.) Hierarchical clustering tree based ongene-stressor interactions. Closeness of relation between a pair of stressors is quantified by the vertical length of the tree branches separating the stressors.

Several genes had strong interactions in some stressors but not others. For example, knockdown of the Hsp90 co-chaperone Sti1 (Chang et al., 1997; Nicolet and Craig, 1989) had aggravating interactions specifically with macbecin and canavaine, two stressors which have been demonstrated to reduce Hsp90 availability (Alford and Brandman, 2018), and unexpectedly, bortezomib (Figure 5C). Knockdown of GCN3, a gene involved in translation initiation regulation (Yang and Hinnebusch, 1996), had the strongest aggravating interactions in the screen, hyperinducing the response to hydrogen peroxide, amino acid starvation, and arsenite stress. Knockdown of Snf4, a Snf1 kinase subunit that plays a key role in glucose-repressed gene transcription (McCartney and Schmidt, 2001), had a strong aggravating interaction only with glucose starvation. Similar to what we observed in the heat shock time course, we saw distinct interaction profiles from the chaperone genes *SSA1* and *SSA2*, with the *SSA1* knockdown having generally weaker blocking interactions, as well as aggravating interactions with AZC and canavanine, two stressors which were strongly blocked by *SSA2* knockdown.

To systematically evaluate which classes of genes are important for the HSR under different conditions, we analyzed which GO categories had strong interaction scores with each of the stress conditions (Figure 5D,E). We observed a wide diversity of stress-specific interactions. For example knockdown of translation-related and mitochondria-related GO categories aggravated the hydrogen peroxide, ethanol, and glucose starvation responses, while blocking the arsenite response. Additionally, proteasome knockdown significantly blocked most responses, but had an aggravating interaction with macbecin and bortezomib. Enrichment scores for all GO categories are provided in table S2. The diversity of genetic interactions observed between stressors and the HSR likely reflects effects of the knockdowns on the damage caused by the stressor as well as on stress-specific signaling to the HSR.

A prevalent model for HSR regulation is that the HSR senses changes in chaperone availability, becoming more active when chaperones become less available. In yeast, the Hsp70 and Hsp90 chaperones have both been proposed to have this role. Thus, lowering levels of either of these chaperones should cause constitutive HSR activation and a loss of sensitivity to stressors (i.e. a blocking interaction). Comparing HSR reporter mRNA induction in each stressor to interaction scores for both *SSA2* and *HSC82* (the gene encoding Hsp90) revealed blocking interaction scores with these chaperones that generally scaled with HSR induction strength, showing that both chaperones were important for responding to stressors (Figure S3A). Yet some interactions were specific to each chaperone. *HSC82*’s interaction with macbecin was stronger than expected based on HSR induction, consistent with macbecin’s inhibition of Hsp90.

Notably, cycloheximide had strong blocking interactions with chaperone genes, even though it does not substantially activate the HSR. This may occur through inhibition of translation, which lowers the load of misfolded proteins in the cell and thus may block the HSR induction that occurs when chaperones are downregulated. Strikingly, plotting basal induction vs cycloheximide interactions revealed a negative correlation (correlation coefficient =-0.23), similar to the effect seen with heat shock (Figure S3B). Hierarchical clustering of stressors revealed that cycloheximide clustered with late heat shock (40 and 120 min), and was more similar to these stressors than were early heat shock (10 and 20 min) (Figure 5F). Thus, most genetic perturbations that affect the HSR are dependent upon translation, consistent with reports that the bulk of misfolded proteins in a cell are newly translated, nascent chains (Medicherla and Goldberg, 2008). Translation-dependent HSR activation may explain blocking interactions observed in other stressors with low HSR induction, such as DTT, tunicamycin, and nutrient starvation.

### Gcn3-dependent translation of the HSR reporter

The knockdown with the strongest and most specific stress-gene interactions was the translation initiation factor, *GCN3*, displaying strong aggravating interactions with hydrogen peroxide, amino acid starvation, and arsenite (Figure 5C). *GCN3* is a component of eIF2B, the nucleotide exchange factor for eIF2. *GCN3* has been shown to sense eIF2 phosphorylation, and prevent nucleotide exchange of eIF2 thereby halting translation (Yang and Hinnebusch, 1996). While each GCN3 interacting stressor has been reported to cause eIF2 phosphorylation (Clemens, 2001), other stressors which did not interact with the *GCN3* knockdown such as heat shock, glucose starvation, tunicamycin, and DTT also result in eIF2 phosphorylation (Clemens, 2001). Phosphorylation of eIF2 is likely important for the role of *GCN3* in the observed stress-specific regulation of the HSR because knockdown of the kinase which phosphorylates eIF2, *GCN2*, shows a similar interaction with the same stressors (Figure 5C).

To further study the effects of *GCN3* knockdown on the HSR, we used a single CRISPR sgRNA selected from the pool targeting *GCN3*. The HSR reporter encodes GFP, allowing us to measure its output by GFP fluorescence. We measured this fluorescence in a condition that was predicted to be aggravated by *GCN3* knockdown, arsenite, as well as one that should be unaffected, AZC. Surprisingly, we found that the activity of a fluorescent HS reporter was not significantly changed by GCN3 knockdown in arsenite (relative to a “no target” control sgRNA carrying the same barcode) rather than increasing as expected (Figure 6A). Because ReporterSeq measures mRNA levels (not protein), we measured mRNA levels under the same conditions via qPCR and observed an 8 fold increase from the *GCN3* knockdown in arsenite relative to control (Figure 6B). As expected, the qPCR experiment showed no substantial effect of *GCN3* knockdown on AZC mRNA levels. This suggests that *GCN3* increases translation efficiency of the HSR reporter under arsenite stress.

**Figure 6.**
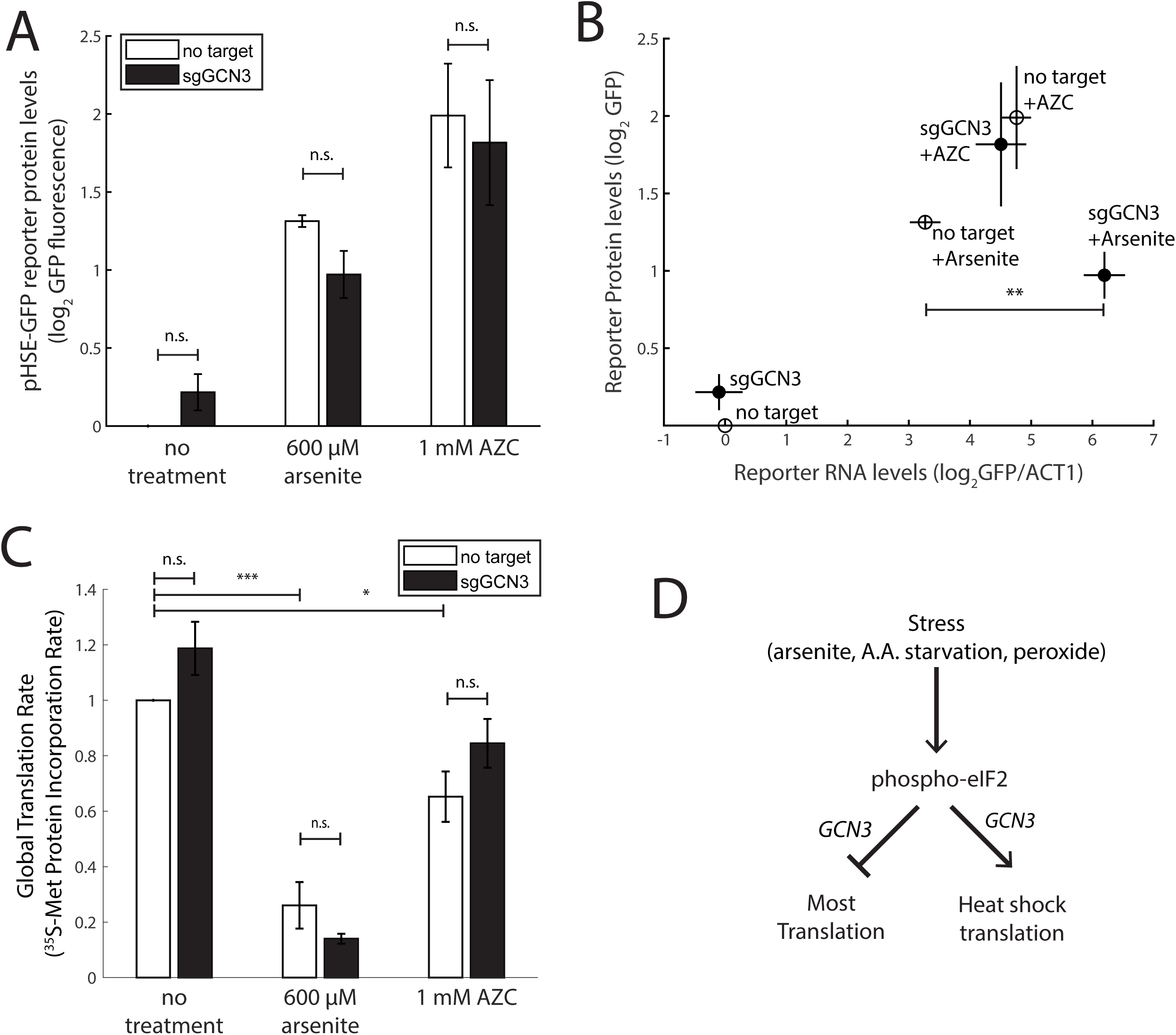
GCN3 is required for the efficient translation of the HSR reporter under arsenite stress. A.) GFP fluorescence of the HSR reporter under the indicated stress conditions and CRISPRi targeting either no gene or *GCN3*. B.) mRNA levels (relative to *ACT1* mRNA) versus protein levels of the HSR reporter as measured by qPCR and GFP fluorescence with the indicated conditions and sgRNAs. C.) Global translation rate as measured by ^35^S-Met incorporation into total protein for the indicated sgRNAs and conditions. D.) Summary diagram *GCN3*’s effect on translation efficiency in selected stressors. All results are normalized to the “no target” sgRNA, unstressed condition. Error bars are standard errors of at least 3 zindependent biological replicates. P-values are indicated for select pairs of conditions based on the following key: n.s.-p>0.05; *-p<0.05; **-p<0.01; ***-p<0.001; ****-p<0.0001.

To determine if GCN3-dependent translation in arsenite was a general phenomenon or specific to the HSR reporter, we measured global translation rates in arsenite and AZC stress, with or without *GCN3* knockdown via incorporation of ^35^S-methionine (Figure 6C). Both stressors decreased translation rates, with AZC causing a ∼30% reduction relative to wildtype and arsenite causing a ∼70% reduction. *GCN3* knockdown, however, caused no significant change in the translation rate in any of the three conditions. We thus conclude that the effect of *GCN3* knockdown on the translation of the HSR reporter in arsenite is not reflective of global changes in translation rate and is likely specific to the HSR reporter (Figure 6D).

## Discussion

We dissected the genetic modulators of the HSR using ReporterSeq, a pooled, high throughput screening method that measures the effect of genome-wide genetic perturbations on the expression of a single reporter-driven RNA. ReporterSeq couples gene expression to levels of barcodes that can be read with deep sequencing, thus enabling pooled, full genome screening without the need for cell enrichment (e.g. via fluorescence-based cell sorting). This allowed us to identify known and novel regulators of the HSR and measure their roles in responding to diverse stressors in a time-resolved manner. Because ReporterSeq measures mRNA levels, we were able to observe modulators that specifically changed mRNA levels and not protein levels, such as *GCN3*, which we discovered increases translation efficiency of the heat shock reporter in some stress conditions.

Our implementation of ReporterSeq featured multiple full-genome screens that can be completed on a single day and read out using a single sequencing run. We expect ReporterSeq results to improve with greater depth of sequencing, and sample size, and replicates. Yet even with the relatively low level of sequencing in this study (∼4-10M reads per sample; table S6), we identified myriad stress-specific regulators of the HSR, including known and novel regulators, creating the most comprehensive genetic view of any protein quality control pathway to date. The scale and precision of ReporterSeq is only limited by growth conditions and DNA sequencing, not cell enrichment, microfluidics, or cell processing.

The strongest HSR-regulating pathway identified by our screen was the Hsp70/Hsp40 pair, *SSA2*/*SIS1*. Knockdown of *SIS1* or *SSA2* desensitized cells to every stressor except for the proteasome inhibitor bortezomib, a weak activator of the HSR. This is consistent with studies demonstrating direct binding between Hsp70 and Hsf1 (Zheng et al., 2016) and suggests that Sis1 is the primary Hsp40 facilitating this interaction. The blocking interactions between stressors and SSA2/SIS1 could be a result of all signaling to Hsf1 being transduced through SSA2/SIS1 or simply that this perturbation is so strong that it saturates the reporter and prevents other means of activating the HSR. We identified one strong regulator of the HSR, Asc1, as an HSR regulator independent of Ssa2/Sis1. Future studies may allow a more systematic delineation of the independence of pathways we identified as driving the HSR.

Both the Hsp90 inhibitor, macbecin, and the proteasome inhibitor, bortezomib, had somewhat similar global gene interaction profiles that differed from other stressors. They both featured aggravating interactions with the proteasome, Hsp90 binding genes, and *SSA1*. Furthermore, bortezomib was the only stressor to positively interact with *SIS1* and *SSA2* while macbecin had little or no interaction with these genes. This suggests that macbecin and bortezomib may activate the HSR via a separate mechanism from SSA2/SIS1, and is consistent with a previous study proposing separate activation mechanisms of the HSR by Hsp70 (*SSA2*) and Hsp90 (Alford and Brandman, 2018).

We observed distinct regulators of the early and late response to heat. This likely arises from separate challenges cells face at each phase of heat shock. In the immediate response to heat, a large fraction of proteins misfold due to the mismatch between folding capacity and demand. Regulators that block this early time point likely block sensing the immediate change in misfolded proteins (e.g. Hsf1, Sis1). Regulators of later time points likely are required for cell maintenance at high temperatures (e.g. Hsp42, Btn2 and Opi10).

The mechanisms by which many of the gene and gene categories we identified regulate the HSR are at this moment unknown. For example, a major mystery is how genes regulating translation and mitochondria, which showed strong interactions with multiple conditions, regulate the HSR. Additionally, how the ras/cAMP pathway and TAZ1, which has been proposed to be essential for protein homeostasis (de Taffin de Tilques et al., 2018), regulate the HSR is poorly understood. Interaction data from our ReporterSeq experiments may be useful in formulating hypotheses that can be tested with detailed follow-up experiments.

All of the discoveries in this work were made possible by the observational power afforded by ReporterSeq. Its full genome, multi stress, time-resolved view of stress-gene interactions allowed the most comprehensive view of the HSR to date. Furthermore, because ReporterSeq measures only mRNA levels, stressors that blocked translation could be studied. Because many stressors block translation, ReporterSeq is ideally suited to study many stress pathways which have been difficult to comprehensively evaluate because they are co-regulated with translation and thus fluorescence- and luciferase-based assays that require translation are limited in effectiveness. Indeed, contemporaneous to this work, a similar technique was used to dissect the genetic regulators of Gcn4(Muller et al., 2020), regulator of the yeast general amino acid control response, a stress-response homologous to the integrated stress response in metazoans. Applied broadly, ReporterSeq may significantly advance the study of cellular pathways *in vivo*.

## Acknowledgements

We thank L. Persson, J. Work, C. Sitron, J. Giafaglione, J. Park, Drs. Z. Davis, Z. Jaafar, D. Khan, R. Rohatgi, M. Krasnow, J. Weissman, L. Gilbert, and C. Jan for helpful suggestions throughout the project and in preparation of the manuscript. Funding was provided by the Lucille P. Markey Basic Biomedical Research Fellowship and NIH 5 T32 GM007276 to BDA, and NSF Awards 1813049 and 1704417 and ONR YIP award N00014-18-1-2295 to GV, and R01GM115968 to OB.

## Methods

### Yeast strains and growth conditions

All experiments were performed with BY4741 yeast. The *ssa2* knockout (yJP385) was made using *NAT* replacement of the *SSA2* ORF, and selected with nourseothricin. Yeast were grown at 30°C (unless otherwise indicated) in synthetic defined (SD) media with the appropriate dropout. SD media used for growth of yeast cultures contained: 2% w/v dextrose (Thermo Fisher Scientific, Waltham, MA), 13.4 g/L Yeast Nitrogen Base without Amino Acids (BD Biosciences, San Jose, CA), 0.03 g/L L-isoleucine (Sigma-Aldrich, St. Louis, MO), 0.15 g/L L-valine (Sigma-Aldrich), 0.04 g/L adenine hemisulfate (Sigma-Aldrich), 0.02 g/L L-arginine (Sigma-Aldrich), 0.03 g/L L-lysine (Sigma-Aldrich), 0.05 g/L L-phenylalanine (Sigma-Aldrich), 0.2 g/L L-Threonine (Sigma-Aldrich), 0.03 g/L L-tyrosine (Sigma-Aldrich), 0.018 g/L L-histidine (Sigma-Aldrich), 0.09 g/L L-leucine (Sigma-Aldrich), 0.018 g/L L-methionine (Sigma-Aldrich), g/L L-tryptophan (Sigma-Aldrich), and 0.018 g/L uracil (Sigma-Aldrich).

### Plasmid library construction

Approximately 12 sgRNAs were designed to target the 5’ end of each protein-coding and non-coding gene in the genome (between -750 and +50 nucleotides from the start of the ORF with priority toward sgRNAs closer to the ORF start) based on the Saccharomyces Genome Database (Cherry et al., 2012). A pool of oligonucleotides was ordered (CustomArray, Inc. Bothell, WA) containing these sequences and constant flanking sequences for the purposes of cloning (Table S5). A 2µ -URA plasmid was constructed from three DNA fragments using HiFi DNA assembly Master Mix (New England Biolabs, Ipswich, MA) for cloning. The first fragment was a diverse sample of inserts amplified via PCR using KAPA HiFi polymerase (Roche Basel, Switzerland), producing a sequence which contained random barcodes associated with a random sgRNAs from the library (with the appropriate reporter plasmid sequence connecting the two, as well as flanking sequences to enable Gibson cloning). The second fragment was the parent plasmid digested at two locations using the FastDigest enzymes BglI and BglII (Thermo Fisher, Sunnyvale, CA). The final fragment was another region of the plasmid amplified via PCR, which was necessary to connect the previous two pieces. These three fragments were assembled in a 100 µL reaction containing 700 ng of the first fragment, 600 ng of the second fragment, and 400 ng of the final fragment. After a 1 hour incubation at 50C, the reaction was concentrated to 20 µL using a DNA clean and concentrate column (Zymo Research Irvine, CA) and electroporated in 5 separate cuvettes containing 4 µL of the plasmid and 40 µL of ElectroTen-Blue electrocompetent cells (Agilent Santa Clara, CA). After a 1 hour outgrowth in LB at 37C, the plasmid was then grown in 1L of LB containing carbenicillin for 2-3 days until it reached saturation. The plasmid was then purified via maxiprep (Qiagen Hilden, Germany).

### Plasmid library sequencing

Two µg of plasmid was cut with the FastDigest PstI restriction enzyme (Thermo Fisher) and the 900 bp piece was gel purified using a gel DNA extraction kit (Zymo). This piece was then recirculized in a 20 µL reaction containing 60 ng of gel purified DNA and 1 µL T4 DNA Ligase (Thermo Fisher) in buffer. Cutting and recircularizing was necessary to increase efficiency of later sequencing steps for unknown reasons. The ligation product was then amplified with primers containing the appropriate overhangs for high throughput sequencing and then gel purified. This piece was then sequenced through paired end sequencing using custom primers on an Illumina (San Diego, CA) sequencing machine (Miseq or Nextseq).

### Yeast library construction

Yeast were first transformed with a 2u pMET17-dCas9-MxiI/*LEU2* plasmid (adapted from (Gilbert et al., 2013)) using standard methods. Five 5 mL of this yeast was grown overnight in SD -Leu media to saturation. This yeast was then back diluted into a 1L YPD culture to approximately ∼0.1 OD660. This culture was then grown at 30°C with shaking until the OD660 reached ∼0.6. The yeast were transformed with 600 ng of the library plasmid using standard procedures. However, instead of plating the yeast after the heat shock, they were grown in 1L of SD -Ura -Leu for 3 days. The yeast were then either rediluted for experiments or frozen in 1 mL aliquots in media with 30% glycerol at -80°C for later use.

### Yeast sample preparation and screening

Yeast from a saturated library sample were grown overnight to log phase in SD -Ura/Leu. These yeast were then split into a number of cultures of at least 10 mL. Yeast were then either left untreated or stressed under the various conditions for the indicated times (Table S3). The yeast were then vacuum filtered onto nitrocellulose paper and flash frozen in liquid nitrogen.

### Nucleic acid extraction and sequencing sample preparation

RNA was extracted through acid-phenol extraction. The frozen yeast pellets were first resuspended in 2.7 mL AE Buffer (50 mM sodium acetate pH 5.5, 10 mM EDTA), 0.3 mL 10% SDS, and 3 mL acid phenol:chloroform (Thermo Fisher). This mixture was heated to 65°C and shaken at 1400 rpm for 10 minutes followed by 5 minutes on ice. The solution was then spun at 13,000g for 15 minutes. The supernatant was then added to 3 mL of chloroform and spun at 15,000g for 5 minutes. The supernatant was then purified through a standard isopropanol extraction.

We then used NextFlex oligo-dT beads (PerkinElmer Waltham, MA) to select polyadenylated mRNA from 100 ug of the extracted RNA. This sample was then subjected to DNAse treatment using Turbo DNase (Thermo Fisher) treatment and then purified using AMPure XP beads (Beckman Coulter, Indianapolis, IN). The resulting RNA was reverse transcribed using Multiscribe reverse transcriptase and a gene-specific primer (Thermo Fisher), and the cDNA was purified using alkaline lysis of the RNA followed by another round of AMPure bead purification.

To obtain the yeast DNA samples, frozen yeast was thawed and miniprepped using a Zymoprep Yeast Plasmid Miniprep Kit (Zymo Research).

The purified cDNA and the purified DNA were then amplified at the barcode region using primers with the appropriate Illumina sequencing ends and indexing barcodes to differentiate each sample. These resulting PCR products were then gel purified, mixed proportionately with each other, and then sequenced using a Illumina next generation sequencing machine (MiSeq, NextSeq, or NovaSeq). The total number of barcode reads matching sgRNAs for each condition is listed in table S6.

### Scoring/statistical analysis

Basal scores and interactions scores were analyzed as pairs of conditions. For the case of the basal score, RNA levels and DNA levels were compared, whereas for the gene-stressors interaction score, stressor RNA was compared to unstressed RNA. When an sgRNA mapped to two adjacent genes, both genes were assigned the sgRNA. For each pair of conditions, *A* and *B*, and each barcode, *x*, we assign a score, *score*(*x, A, B*), which captures how extreme (positive or negative) the read count is for barcode *x* in condition *A* versus condition *B*. These scores allow us to compare the statistical significance of discrepancies between counts in different conditions, enabling a convenient aggregation to form a score for each gene, or gene cluster. Crucially, the scores take into account 1) the higher relative amount of noise in small counts versus larger counts, 2) the fact that some conditions yield higher read counts across most barcodes.

We now describe and motivate our definition of *score*(*x, A, B*). Let *A*(*x*) denote the logarithm of the count of barcode *x* under condition *A*, and let *B*(*x*) denote the logarithms of the counts for barcode *x* under conditions *B*, scaled by a factor *c*_*A,B*_ that compensates for the possibility that some conditions yield systematically higher counts than others. After this scaling, for most barcodes, *x*′, it is the case that *A*(*x*′) ≈ *B*(*x*′).

Let 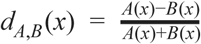 denote the normalized discrepancy in these values, and note that this quantity is bounded between ± 1. The score is this discrepancy, *d*_*A,B*_ (*x*) after the median value of this is subtracted, normalized by a factor *S*_*A*(*x*),*B*(*x*)_ that takes into account how much variance is typical in this quantity, given the values of *A*(*x*) and *B*(*x*). Note that we expect *d*_*A,B*_ (*x*) to have a larger variance due to random noise in the experiment and measurement when *A*(*x*) and *B*(*x*) correspond to small counts. This scaling factor *S*_*A*(*x*),*B*(*x*)_ is defined to be an outlier-robust analog of the standard deviation of *d*_*A,B*_ (*x*′) computed across the set of barcodes *x*′ for which *A*(*x*′) + *B*(*x*′) ≈ *A*(*x*) + *B*(*x*).

The following pseudocode formalizes the above calculation of the scores for each barcode in each pair of conditions:

### Calculate barcode score

**Input:** Barcode *x*, read counts for all barcodes in conditions *A* and *B*, and scaling factor *c(A,B)*.

**Output:** *score*(*x,A,B*)

- Define parameters Cohort_size = 10,000 and Clip_Param = 100.
- Let *A*(*x*) denote the logarithm of the read count for barcode *x* in condition *A*, and let *B*(x) denote the logarithm of the read count for barcode *x* in condition *B* scaled by a constant *c(A,B)* defined as the ratio of the median of the logarithms of the top 100 counts in condition *A*, to the median of the logarithms of the to 100 counts in condition *B*.
- Define the set of barcodes *X* = {x’ : |A(x’)+B(x’) - A(x)+B(x)| < delta},
where delta is the smallest value such that |*X*| ≥ Cohort_size.
- For each *x’* in *X*, let *d*(x’) = (A(x)-B(x))/(A(x)+B(x))
- Define *S* = truncatedStandardDeviation(d(X),Clip_Param), which is the standard deviation of the multiset of values of d(x’) taken across all elements x’ of X, after the largest and smallest Clip_Param values are removed. (E.g. if Clip_Param = 100, the largest and smallest 100 values are removed prior to computing the standard deviation.)
- Return *score(x,A,B)*=[d(x)-median(d(X))]/S.

Gene Ontology (GO) scores were calculated based on the above calculated scores after saturation of the top and bottom 0.5% scores (to prevent strong outliers from dominating a category score). For each GO category, the distribution of scores for genes within that category and those of the entire distribution (scores for all genes) were compared using a Student’s t-test. The GO score was the natural log of the reciprocal of the resultant p-value from the Student’s t-test. The GO score sign indicates in whether the mean of the gene scores for that gene category was above or below zero.

Hierarchical clustering was performed using Cluster 3.0(de Hoon et al., 2004), using uncentered correlation similarity metric and complete linkage clustering method. Clustering was performed on all genes except tRNAs and retrotransposons.

### Flow cytometry

Yeast were grown overnight from a saturated culture to log phase (OD < 0.6), such that they were growing in log phase for at least 14 hours. These yeast were then measured on an Accuri flow cytometer at 10,000 yeast per sample, under the fast flow setting.

### Global translation rate measurement

Yeast were grown overnight to an OD of ∼0.1 subjected to the indicated stressed or unstressed condition for one hour. Subsequently, ^35^S-methionine was added to each culture and samples were taken at the 0 minute, 20 minute, and 1 hour time points. These 1 ml samples were centrifuged at 13,000 RPM for 30 seconds and the resultant pellets of yeast were frozen in liquid nitrogen. Yeast were then lysed and protein was extracted via TCA precipitation. The TCA precipitation involved first incubating the yeast pellet in 240 µL alkaline buffer (750 µM NaOH,7.5% β-mercaptoethanol) for 10 minutes on ice. Then, 240 µL chilled 50% TCA was added, the mixture was vortexed, and then it was incubated for another 10 minutes on ice. It was spun for 10 minutes at 13,000 rpm and then the supernatant was removed. The pellet was then incubated with 500 µL of 90% acetone for 20 minutes at -20°C. The sample was then spun at 13,000 rpm for another 10 minutes and then the acetone was removed and the pellet was allowed to air dry. The pellet was then resuspended in a 100 µL resuspension buffer (50 mM Tris-Cl pH 7.5, 50 mM NaCl, 2.5% SDS). Total protein amount was measured from 10 µL of the sample using the Pierce BCA protein assay kit (Thermo Fisher). The levels of radioactive methionine were measured by scintillation counting with 10-80 µL of the resuspended protein. The scintillation counting was done at multiple concentrations of protein to ensure that the count was in the linear range of the machine.

### Quantitative PCR (qPCR)

RNA was extracted and DNase treated according to the protocol described above. One µg of RNA was then reverse transcribed using Multiscribe reverse transcriptase (Thermo Fisher) and random hexamers in a 25 µL reaction. One µL of the cDNA was then mixed with Luna qPCR master mix (New England Biolabs) and primers (to a final concentration 100 nM) which amplified either GFP or *ACT1*, the reference gene. These mixtures were measured in triplicate for each sample-primer combination with a ViiA 7 qPCR machine (Thermo Fisher)

---

**Figure S1.**
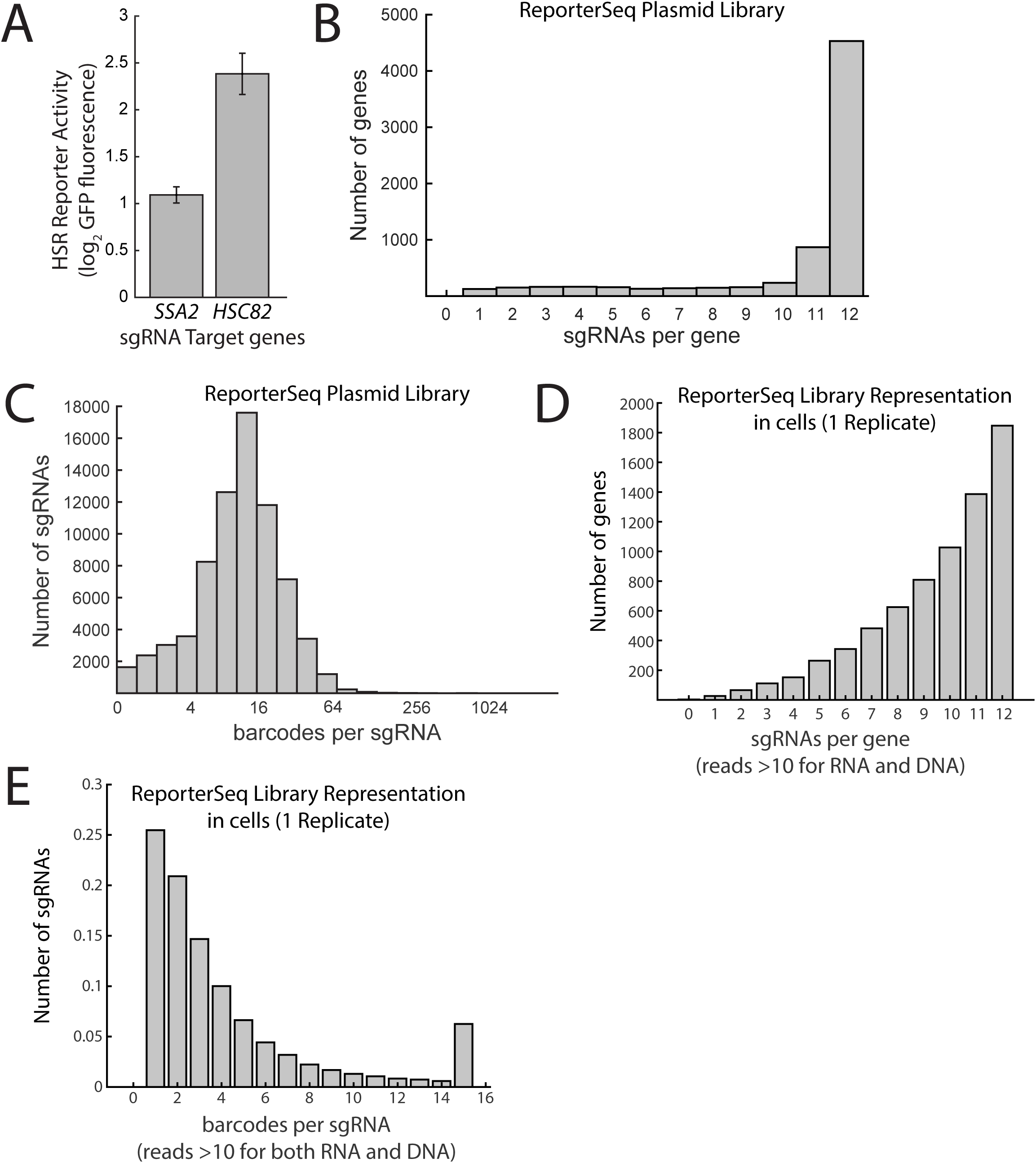
Diversity of ReporterSeq Library. A.) GFP fluorescence of yeast containing the pHSE-GFP synthetic Hsf1 reporter with the sgRNAs targeting the indicated genes. The fluorescence is relative to a non-targeting sgRNA. Error bars are standard errors of at least 3 replicates. B.) Histogram of the number of sgRNAs targeting each gene found in the plasmid library through paired-end sequencing. C.) Histogram of the number of barcodes associated with each sgRNA found in the library through paired end sequencing. The y-axis is a log base-2 scale of barcode counts. D) Histogram of sgRNAs per gene in transfected cells with both DNA and RNA counts 10 or above. E) Histogram of barcodes per sgRNA with both DNA and RNA counts 10 or above.

**Figure S2.**
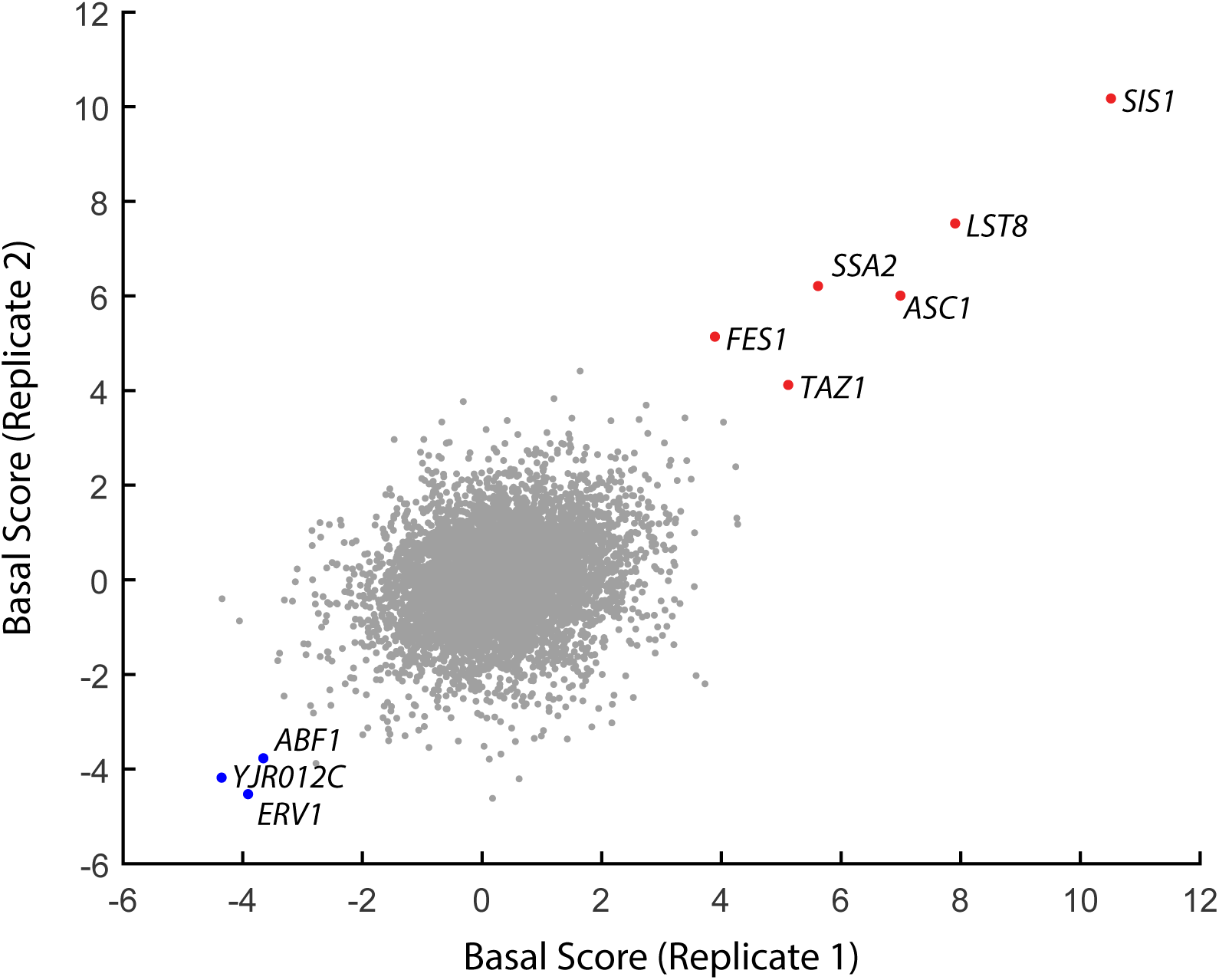
Comparison of two independent biological replicates of ReporterSeq measuring basal HSR regulation. Replicates are from two separate yeast transformations with the ReporterSeq plasmid library.

**Figure S3.**
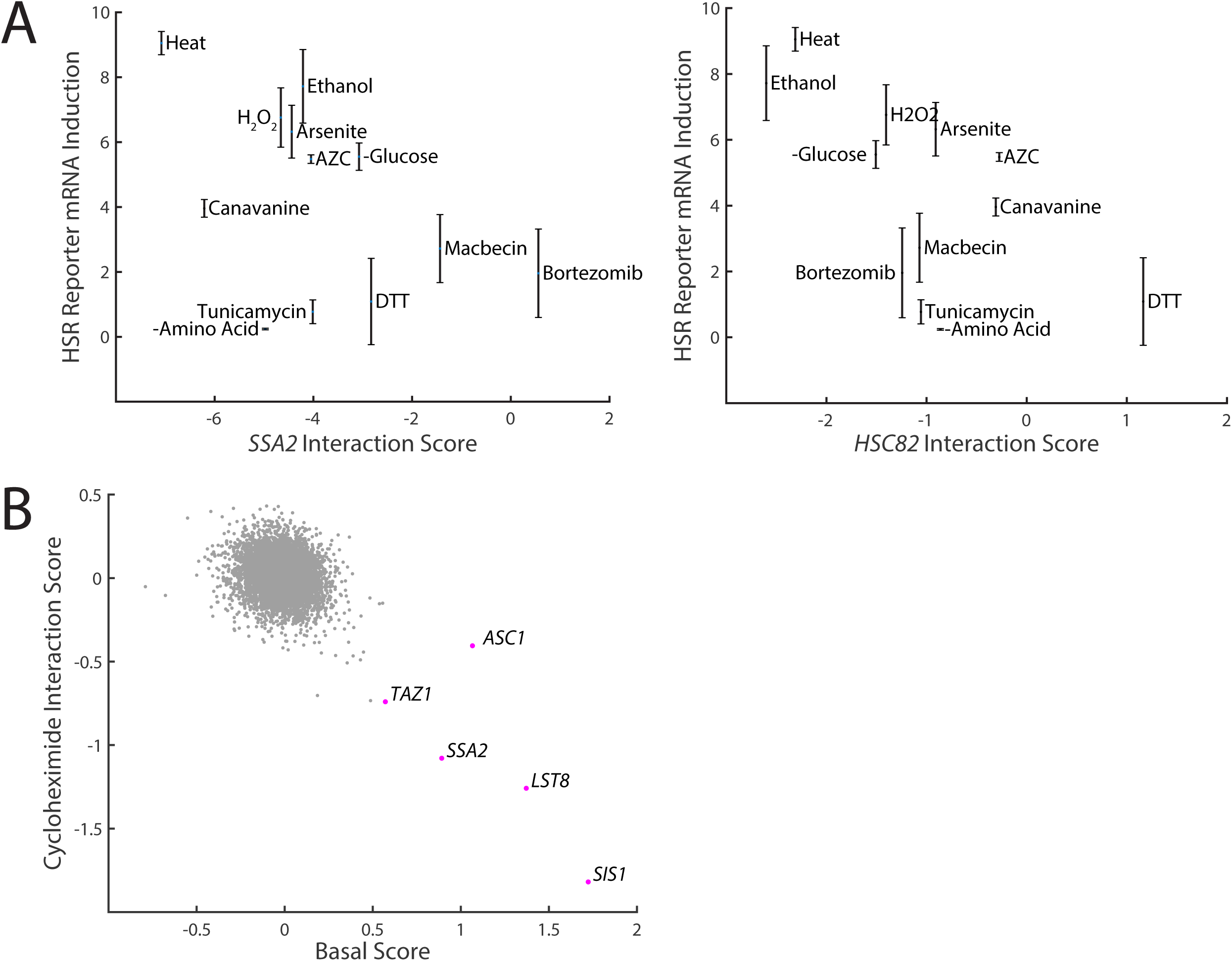
Comparisons of proteotoxic stressor effects on the HSR. A.) HSR reporter mRNA induction versus the interaction scores of *SSA2* (left panel) and *HSC82* (right panel) with each stressor. B.) Basal score versus cycloheximide interaction score for each gene. Genes with a high basal score are labeled.

## References

Akerfelt M, Morimoto RI, Sistonen L. 2010. Heat shock factors: integrators of cell stress, development and lifespan. Nat Rev Mol Cell Biol 11 :545–555.

Alford BD, Brandman O. 2018. Quantification of Hsp90 availability reveals differential coupling to the heat shock response. J Cell Biol 217 :3809–3816.

Ali A, Bharadwaj S, O’Carroll R, Ovsenek N. 1998. HSP90 interacts with and regulates the activity of heat shock factor 1 in Xenopus oocytes. Mol Cell Biol 18 :4949–4960.

Anckar J, Sistonen L. 2011. Regulation of HSF1 function in the heat stress response: implications in aging and disease. Annu Rev Biochem 80 :1089–1115.

Ashburner M, Ball CA, Blake JA, Botstein D, Butler H, Cherry JM, Davis AP, Dolinski K, Dwight SS, Eppig JT, Harris MA, Hill DP, Issel-Tarver L, Kasarskis A, Lewis S, Matese JC, Richardson JE, Ringwald M, Rubin GM, Sherlock G. 2000. Gene ontology: tool for the unification of biology. The Gene Ontology Consortium. Nat Genet 25 :25–29.

Brandman O, Stewart-Ornstein J, Wong D, Larson A, Williams CC, Li G-W, Zhou S, King D, Shen PS, Weibezahn J, Dunn JG, Rouskin S, Inada T, Frost A, Weissman JS. 2012. A ribosome-bound quality control complex triggers degradation of nascent peptides and signals translation stress. Cell 151 :1042–1054.

Campanella C, Pace A, Caruso Bavisotto C, Marzullo P, Marino Gammazza A, Buscemi S, Palumbo Piccionello A. 2018. Heat Shock Proteins in Alzheimer’s Disease: Role and Targeting. Int J Mol Sci b>19. doi: 10.3390/ijms19092603

Chang HC, Nathan DF, Lindquist S. 1997. In vivo analysis of the Hsp90 cochaperone Sti1 (p60). Mol Cell Biol 17 :318–325.

Cherry JM, Hong EL, Amundsen C, Balakrishnan R, Binkley G, Chan ET, Christie KR, Costanzo MC, Dwight SS, Engel SR, Fisk DG, Hirschman JE, Hitz BC, Karra K, Krieger CJ, Miyasato SR, Nash RS, Park J, Skrzypek MS, Simison M, Weng S, Wong ED. 2012. Saccharomyces Genome Database: the genomics resource of budding yeast. Nucleic Acids Res 40 :D700–5.

Clemens MJ. 2001. Initiation factor eIF2 alpha phosphorylation in stress responses and apoptosis. Prog Mol Subcell Biol 27 :57–89.

Collins SR, Miller KM, Maas NL, Roguev A, Fillingham J, Chu CS, Schuldiner M, Gebbia M, Recht J, Shales M, Ding H, Xu H, Han J, Ingvarsdottir K, Cheng B, Andrews B, Boone C, Berger SL, Hieter P, Zhang Z, Brown GW, Ingles CJ, Emili A, Allis CD, Toczyski DP, Weissman JS, Greenblatt JF, Krogan NJ. 2007. Functional dissection of protein complexes involved in yeast chromosome biology using a genetic interaction map. Nature 446 :806–810.

Conlin LK, Nelson HCM. 2007. The natural osmolyte trehalose is a positive regulator of the heat-induced activity of yeast heat shock transcription factor. Mol Cell Biol 27 :1505–1515.

de Hoon MJL, Imoto S, Nolan J, Miyano S. 2004. Open source clustering software. Bioinformatics 20 :1453–1454.

de Taffin de Tilques M, Lasserre J-P, Godard F, Sardin E, Bouhier M, Le Guedard M, Kucharczyk R, Petit PX, Testet E, di Rago J-P, Tribouillard-Tanvier D. 2018. Decreasing cytosolic translation is beneficial to yeast and human Tafazzin-deficient cells. Microb Cell Fact 5 :220–232.

Gilbert LA, Larson MH, Morsut L, Liu Z, Brar GA, Torres SE, Stern-Ginossar N, Brandman O, Whitehead EH, Doudna JA, Lim WA, Weissman JS, Qi LS. 2013. CRISPR-mediated modular RNA-guided regulation of transcription in eukaryotes. Cell 154 :442–451.

Guarente L, Mason T. 1983. Heme regulates transcription of the CYC1 gene of S. cerevisiae via an upstream activation site. Cell 32 :1279–1286.

Hahn J-S, Thiele DJ. 2004. Activation of the Saccharomyces cerevisiae heat shock transcription factor under glucose starvation conditions by Snf1 protein kinase. J Biol Chem 279 :5169–5176.

Haitani Y, Shimoi H, Takagi H. 2006. Rsp5 regulates expression of stress proteins via post-translational modification of Hsf1 and Msn4 in Saccharomyces cerevisiae. FEBS Lett 580 :3433–3438.

Haitani Y, Takagi H. 2008. Rsp5 is required for the nuclear export of mRNA of HSF1 and MSN2/4 under stress conditions in Saccharomyces cerevisiae. Genes Cells 13 :105–116.

Haslbeck M, Braun N, Stromer T, Richter B, Model N, Weinkauf S, Buchner J. 2004. Hsp42 is the general small heat shock protein in the cytosol of Saccharomyces cerevisiae. EMBO J 23 :638–649.

Heikkinen HL. 2003. Initiation-mediated mRNA decay in yeast affects heat-shock mRNAs, and works through decapping and 5’-to-3’ hydrolysis. Nucleic Acids Research. doi: 10.1093/nar/gkg474

Jacobson T, Navarrete C, Sharma SK, Sideri TC, Ibstedt S, Priya S, Grant CM, Christen P, Goloubinoff P, Tamás MJ. 2012. Arsenite interferes with protein folding and triggers formation of protein aggregates in yeast. J Cell Sci 125 :5073–5083.

Kose S, Furuta M, Imamoto N. 2012. Hikeshi, a nuclear import carrier for Hsp70s, protects cells from heat shock-induced nuclear damage. Cell 149 :578–589.

Krakowiak J, Zheng X, Patel N, Feder ZA, Anandhakumar J, Valerius K, Gross DS, Khalil AS, Pincus D. 2018. Hsf1 and Hsp70 constitute a two-component feedback loop that regulates the yeast heat shock response. Elife 7. doi: 10.7554/eLife.31668

Lee P, Cho B-R, Joo H-S, Hahn J-S. 2008. Yeast Yak1 kinase, a bridge between PKA and stress-responsive transcription factors, Hsf1 and Msn2/Msn4. Molecular Microbiology. doi: 10.1111/j.1365-2958.2008.06450.x

Lee P, Kim MS, Paik S-M, Choi S-H, Cho B-R, Hahn J-S. 2013. Rim15-dependent activation of Hsf1 and Msn2/4 transcription factors by direct phosphorylation inSaccharomyces cerevisiae. FEBS Letters. doi: 10.1016/j.febslet.2013.10.004

Lindquist S. 1986. The Heat-Shock Response. Annual Review of Biochemistry. doi: 10.1146/annurev.bi.55.070186.005443

Malinovska L, Kroschwald S, Munder MC, Richter D, Alberti S. 2012. Molecular chaperones and stress-inducible protein-sorting factors coordinate the spatiotemporal distribution of protein aggregates. Mol Biol Cell 23 :3041–3056.

McCartney RR, Schmidt MC. 2001. Regulation of Snf1 kinase. Activation requires phosphorylation of threonine 210 by an upstream kinase as well as a distinct step mediated by the Snf4 subunit. J Biol Chem 276 :36460–36466.

Medicherla B, Goldberg AL. 2008. Heat shock and oxygen radicals stimulate ubiquitin-dependent degradation mainly of newly synthesized proteins. J Cell Biol 182 :663–673.

Morano KA, Grant CM, Moye-Rowley WS. 2012. The response to heat shock and oxidative stress in Saccharomyces cerevisiae. Genetics 190 :1157–1195.

Morimoto RI. 2011. The Heat Shock Response: Systems Biology of Proteotoxic Stress in Aging and Disease. Cold Spring Harb Symp Quant Biol 76 :91–99.

Muller R, Meacham ZA, Ferguson L, Ingolia N. 2020. CiBER-seq dissects genetic networks by quantitative CRISPRi profiling of expression phenotypes. bioRxiv. doi: 10.1101/2020.03.29.015057

Neckers L, Workman P. 2012. Hsp90 molecular chaperone inhibitors: are we there yet? Clin Cancer Res 18 :64–76.

Neef DW, Jaeger AM, Gomez-Pastor R, Willmund F, Frydman J, Thiele DJ. 2014. A direct regulatory interaction between chaperonin TRiC and stress-responsive transcription factor HSF1. Cell Rep 9 :955–966.

Nicolet CM, Craig EA. 1989. Isolation and characterization of STI1, a stress-inducible gene from Saccharomyces cerevisiae. Mol Cell Biol 9 :3638–3646.

Pincus D, Anandhakumar J, Thiru P, Guertin MJ, Erkine AM, Gross DS. 2018. Genetic and epigenetic determinants establish a continuum of Hsf1 occupancy and activity across the yeast genome. Mol Biol Cell 29 :3168–3182.

Raychaudhuri S, Loew C, Körner R, Pinkert S, Theis M, Hayer-Hartl M, Buchholz F, Hartl FU. 2014. Interplay of acetyltransferase EP300 and the proteasome system in regulating heat shock transcription factor 1. Cell 156 :975–985.

Song J, Kose S, Watanabe A, Son SY, Choi S, Hong H, Yamashita E, Park IY, Imamoto N, Lee SJ. 2015. Structural and functional analysis of Hikeshi, a new nuclear transport receptor of Hsp70s. Acta Crystallogr D Biol Crystallogr 71 :473–483.

Sorger PK. 1990. Yeast heat shock factor contains separable transient and sustained response transcriptional activators. Cell. doi: 10.1016/0092-8674(90)90123-v

Specht S, Miller SBM, Mogk A, Bukau B. 2011. Hsp42 is required for sequestration of protein aggregates into deposition sites in Saccharomyces cerevisiae. J Cell Biol 195 :617–629.

The Gene Ontology Consortium. 2019. The Gene Ontology Resource: 20 years and still GOing strong. Nucleic Acids Research. doi: 10.1093/nar/gky1055

Tong AH, Evangelista M, Parsons AB, Xu H, Bader GD, Pagé N, Robinson M, Raghibizadeh S, Hogue CW, Bussey H, Andrews B, Tyers M, Boone C. 2001. Systematic genetic analysis with ordered arrays of yeast deletion mutants. Science 294 :2364–2368.

Trotter EW, Kao CM-F, Berenfeld L, Botstein D, Petsko GA, Gray JV. 2002. Misfolded proteins are competent to mediate a subset of the responses to heat shock in Saccharomyces cerevisiae. J Biol Chem 277 :44817–44825.

Weids AJ, Ibstedt S, Tamás MJ, Grant CM. 2016. Distinct stress conditions result in aggregation of proteins with similar properties. Sci Rep 6 :24554.

Whitesell L, Lindquist S. 2009. Inhibiting the transcription factor HSF1 as an anticancer strategy. Expert Opin Ther Targets 13 :469–478.

Yang W, Hinnebusch AG. 1996. Identification of a regulatory subcomplex in the guanine nucleotide exchange factor eIF2B that mediates inhibition by phosphorylated eIF2. Mol Cell Biol 16 :6603–6616.

Zheng X, Krakowiak J, Patel N, Beyzavi A, Ezike J, Khalil AS, Pincus D. 2016. Dynamic control of Hsf1 during heat shock by a chaperone switch and phosphorylation. Elife 5. doi: 10.7554/eLife.18638

Zid BM, O’Shea EK. 2014. Promoter sequences direct cytoplasmic localization and translation of mRNAs during starvation in yeast. Nature 514 :117–121.

Zou J, Guo Y, Guettouche T, Smith DF, Voellmy R. 1998. Repression of Heat Shock Transcription Factor HSF1 Activation by HSP90 (HSP90 Complex) that Forms a Stress-Sensitive Complex with HSF1. Cell. doi: 10.1016/s0092-8674(00)81588-3

